# Identification of variation in *Fibromelanosis* region on chromosome 20 for determining the purity of Indonesian Cemani chicken

**DOI:** 10.1101/2022.11.13.516295

**Authors:** Anik Budhi Dharmayanthi, Keiji Kinoshita, Isyana Khaerunnisa, Rona Saumy Safitry, Syam Budi Iryanto, Yohanna, Sutikno, Andi Baso Lompengeng Ishak, M Syamsul Arifin Zein, Yoko Satta, Toyoko Akiyama, Cece Sumantri

## Abstract

Ayam Cemani is a local Indonesian chicken with heavy pigmentation in plumage colour, skin, eyes, and inner body organs. This trait with dermal hyperpigmentation is identical to *Fibromelanosis* (*Fm*) mutation in a Silkie chicken. The causal mutation of the *Fm* trait is due to an inverted duplication and junction of two genomic regions involving the Endothelin3 (*EDN3*) gene on chromosome 20. There are two duplication boundaries; one is specific to the *Fm* allele, the other is common for both *Fm* and *fm*^*+*^ allele. Determining birds that are homozygous or heterozygous at this locus is useful for unifying the *Fm* trait of Cemani populations. This study develops a method for determining the presence or absence of *Fm* mutation by PCR amplification using the inverted sequences specific to the *Fm* allele. Further, it develops the restriction fragment length polymorphism (RFLP) method in regions common to the *Fm* and wild-type *fm*^*+*^ allele. We aim to establish a simple method for detecting homozygous (*Fm*/*Fm*) and heterozygous (*Fm*/*fm*^*+*^) individuals with *Fm* mutation and to clarify the degree of fixation of the *Fm* trait in the Ayam Cemani populations and the association between the phenotype and genotype. The result showed that mostly, the phenotype for Cemani with *Fm*/ *fm*^*+*^ genotype is reddish black in their comb; meanwhile, the Cemani with (*Fm*/*Fm*) genotype showed heavy black pigmentation. Our study concluded that using the PCR-RFLP method. We can discriminate between *Fm* homozygous and heterozygous birds in the Cemani population. Thus, this briefly genotyping method effectively maintains and protects the pure line of Cemani chicken.

## 1. Introduction

Ayam Cemani (black chicken) is a rare local breed on Java Island, Indonesia. The most significant characteristic of Ayam Cemani is hypermelanic pigmentation that covers the entire body, including the plumage, comb, shank, tongue, and eye. Extensive pigmentation can also be found in most of the inner body, such as muscle, intestines, bones, peritoneum and trachea. This unique phenotype is known as *Fibromelanosis* (*Fm*), and Silkie and Ayam Cemani are known as representative breeds carrying the *Fm* mutation.

Based on the Decree of the Minister of Agriculture of the Republic of Indonesia (No: 2487/Kpts/LB/430/8/2012), the Cemani chicken is defined as a particular type of black Kedu chicken with *Fm* mutation. The Kedu chicken is further classified into *Kedu Merah* (red Kedu), *Kedu Hitam* (black Kedu) and *Kedu Putih* (white Kedu) according to the difference in plumage colour, but these do not carry the *Fm* trait (Ismoyowati et al. 2012). Conspicuous characteristics of Cemani chicken are known as dermal hyperpigmentation, which causes a dark colour in internal organs. This phenotype has been associated with the *Fibromelanosis* (*Fm*) locus (Arora et al. 2011, Shinomiya et al. 2012). *Fibromelanosis* is a mutation in chickens indicated by the accumulation of melanin pigment in the internal organs and connective tissue (Hutt, 1949, Muroya et al. 2000, Dorshort et al. 2011). It is known that the Cemani chicken originated from Kedu village, Temanggung city, Central Java. Apart from being known as ornamental chickens from Temanggung, Central Java Province. The origin of the Cemani chicken is still unclear since there is no historical record of the bird. However, Cemani was perhaps selected and bred from Kedu chickens as a variety carrying self-black plumage with *Fm* trait. Further, based on well-known stories passed down from generation to generation, the deep black Cemani chicken has been used for at least hundreds of years for religious and mystical purposes. Some believe cemani chickens have magical powers to repel reinforcements (Sulandari et al. 2007).

Ayam Cemani is currently bred by small farms or individual breeders, primarily in the village of Kedu in Central Java, with some breeders mating with other Kedu chickens and breeds. Some individuals have incomplete Fm traits in recent years, suggesting that the *Fm* traits may not be completely fixed within the population. These changes in the genetic characteristics of Cemani are being threatened, and the problem that the *Fm* trait, which is the most distinguishing feature of Cemani, is not fixed in the Cemani population is emerging.

Estimating the genetic purity for deep-black pigmentation of the Cemani chicken is necessary to conserve the pure-local specific line, Cemani. Although the assumption of the genotype of *Fm* trait of the Cemani has been carried out based on the degree of melanin pigmentation in the beak, feather, skin underwing, and cloaca, the judgment often induces bias results probably due to the effects of other loci associated with pigmentation such as dermal melanin inhibitor (*Id*). Thus, a molecular approach can be used to accurately determine the Fm genotype of Cemani chicken. Previous studies have shown that Fm mutation in Cemani and Silkie chickens is caused by a rearrangement involving inversion and duplication of the genomic region containing *EDN3* on chromosome 20 (Shinomiya et al. 2011, Dorshorst et al. 2011). Dorshorst et al. proposed three duplicate rearrangements. These duplications consist of two duplication regions, duplication region1 (DR1) and duplication region2 (DR2), separated by a 417 kb spacer, and one duplication region is inverted duplication. These rearrangements possess the exact boundaries of 1RD-DR2/2RD-DR1 and DR1-2RD/DR2-1RD. (Figure 1, Dorshorst et al. 2011). Moreover, a recent study in Cemani chicken concluded that FM2 is the best rearrangement for duplication by identifying the patterns and degrees of polymorphism in 417kb spacer (Dharmayanthi et al. 2017).

**Figure 1.**
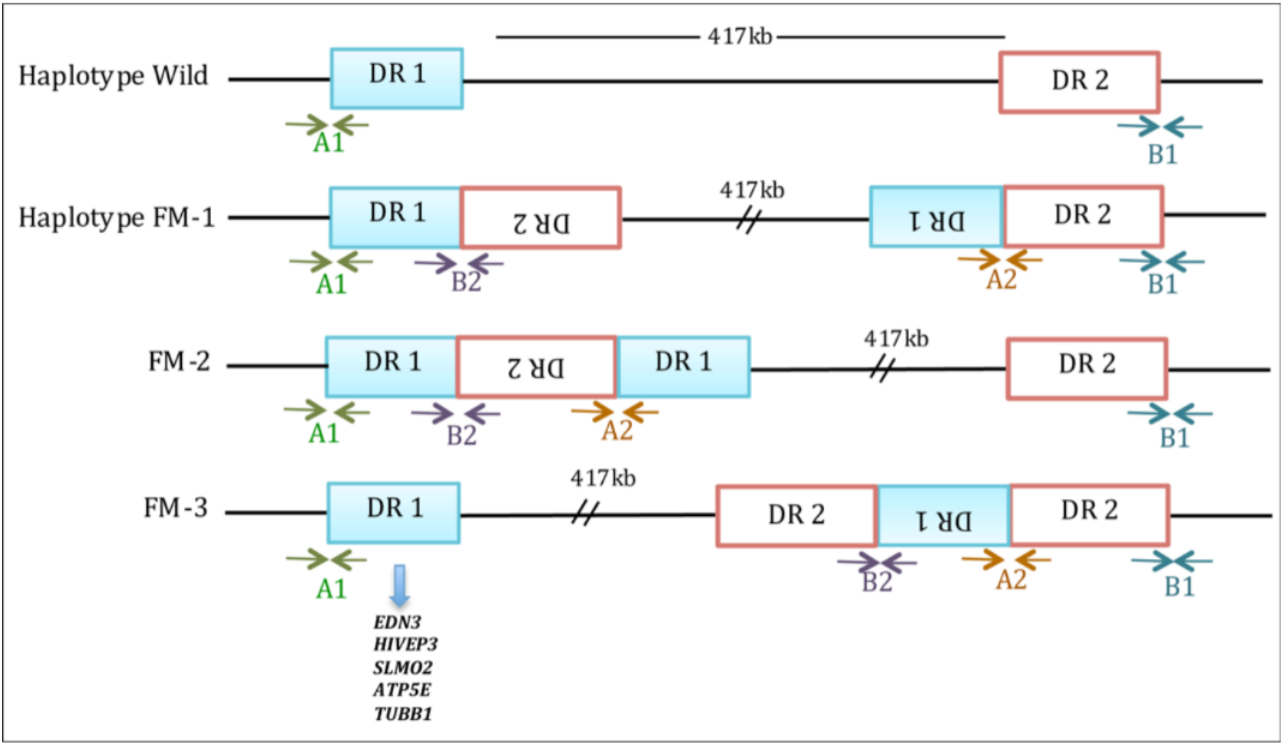
Three possible rearrangements of duplicated DR1 and DR2 in the *Fm* region (Dorshorst et al. 2011 modified in Dharmayanthi et al. 2017).

To date, fixability of the *Fm* mutation trait in the Cemani chicken population based on genotype-phenotype association analysis has not been investigated. Thus, the purpose of the present study is to establish a straightforward molecular method for identifying genotypes of the *Fm* trait in Ayam Cemani populations. We established a PCR-based genotyping method for determining the presence or absence of the *Fm* mutation using primers designed to amplify the *Fm* (haplotype FM-2 in Figure 1) allele-specific region. Furthermore, we established a method for distinguishing birds homozygous and heterozygous for the *Fm* trait by PCR-RFLP or sequencing of SNPs in regions common to *Fm* and wild-type (WT) alleles. The genotype-phenotype association analysis revealed that homozygous and heterozygous genotypes of the *Fm* mutation were associated with the degree of extensive pigmentation of the dermis, especially in the comb and cloaca. This method allows for the selection of breeder chickens for the next generation of Ayam Cemani populations, as well as the complete fixation of the *Fm* trait in the next generation(s). In addition, to avoid genetic inbreeding in the population, the degree of *Fm* fixation introduced into a population can be determined using this method

## 2. Materials and Methods

### 2.1. Chicken sample

In this study, we used a total of 103 samples of Cemani chickens from three different sampling sites; 40 blood samples from the livestock research institute (Balitnak), Bogor; 37 blood samples from Sumber Unggas Indonesia (SUI); and 26 genomic DNA samples (Kedu village, Central Java) preserved in the genetic laboratory, Museum Zoologicum Bogoriense (MZB), Research Center for Biology, Indonesian Institute of Science (RCB-LIPI). Each individual of Cemani was photographed to record its degree of eumelanin pigmentation on parts such as comb, shank, skin underwing and cloaca. Besides, we also used 22 samples of wild-type chickens without *Fm* mutation, consisting of nine Broiler, nine Kedu Putih and four IPB-D1 chickens (Table 5). IPB-D1 chicken is a composite chicken created by interbreeding three Indonesian native chickens (Ayam Pelung, Ayam Sentul, and Ayam Kampung) with Broilers chicken until the fifth generation at Institute Pertanian Bogor (IPB) University.

### 2.2. DNA Extraction

Genomic DNA was extracted using the Geneaid™ DNA Isolation Kit (Blood) following the manufacturing protocol.

### 2.3. PCR amplification and Restriction Fragment Length Polymorphism (RFLP)

Primers for 558 bp specific to the *Fm* allele and 664 bp common to *Fm* and WT alleles were designed based on information from rearrangement FM2 as described above (Figure 1). As the first screening, these two fragments were simultaneously amplified by multiplex PCR using three primers to detect the presence of *Fm*-specific bands and typical bands in 3 populations (Cemani, IPB-D1, Broiler, and Kedu Putih) listed in Table 1. The PCR amplification of each chicken was performed according to the following conditions: the multiplex-PCR was performed in a total volume of 25 μl mix solution containing 12.5μl MyTaq HS Red Mix (Bioline), 40ng of genomic DNA, 10 pmol of each oligonucleotide primer; PCR reaction cycle parameters were denaturation for 1 min at 95°C then 35 cycles of 95°C for 15 sec, 60°C annealing for 15 sec, and 72°C for 10 sec, with a final extension step for 5 min at 72°C. PCR-RFLP genotyping assay was performed as a second screening to distinguish between *Fm* homozygous (*Fm*/*Fm*) and heterozygous (*Fm*/*fm*^*+*^) birds using polymorphism for *Mlu*I restriction enzyme within the standard band of 664 bp. The 664 bp of multiplex-PCR products, including 558 bp fragment, were digested at 37°C overnight with 10 U of *Mlu*I restriction enzyme. The digested products were electrophoresed for 50 minutes at 50V on a 2% agarose gel. Extragen-filtered tips are used in the restriction digest method.

**Table 1.**
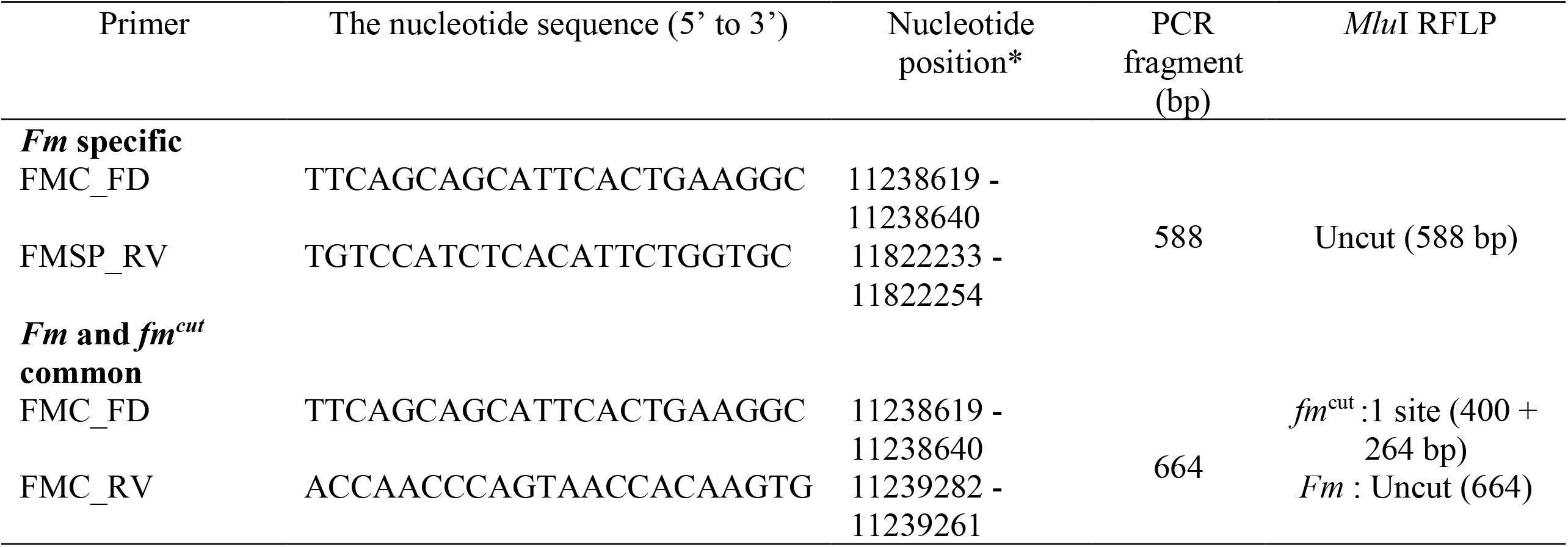
List of primer, primer sequence, position and restriction enzyme.

### 2.4. PCR and Sequencing 664bp fragment length

Since some WT individuals used in this study do not have a *Mlu*I site within 664 bp for PCR-RFLP assay, we performed sequence analysis on individuals with *Fm* mutation that could not distinguish between homozygous and heterozygous and WT individuals with/without *Mlu*I site. The 664bp fragments were amplified with a pair of primers, FMC_RV and FMC_FD, following PCR conditions as multiplex PCR mentioned above (Figure 2). PCR products were then sent to be sequenced in first base Singapore. The sequence data were analysed using Codon Code aligner v9.0.1.

**Figure 2.**
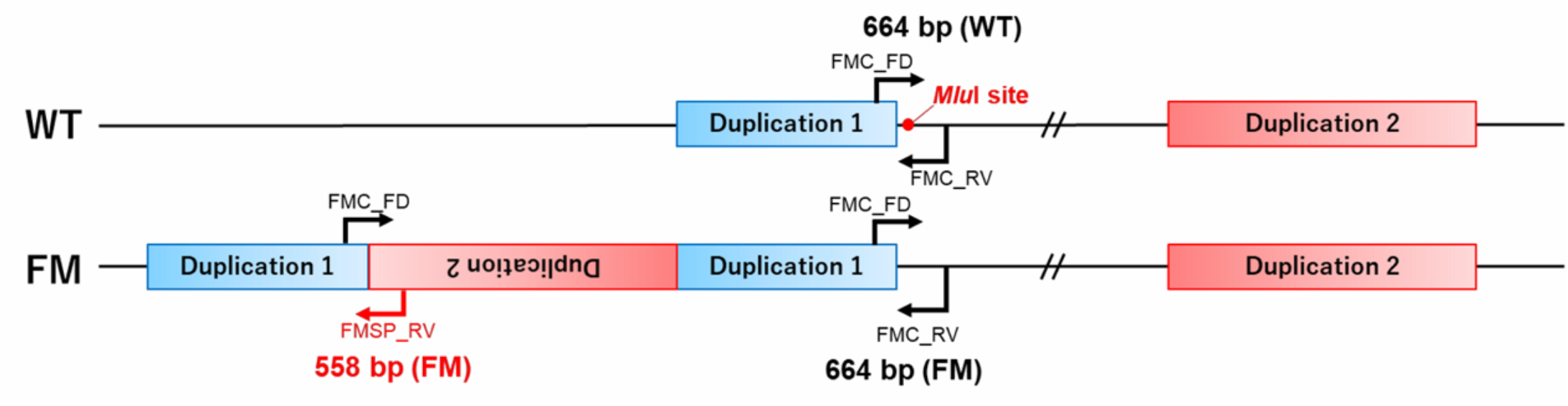
Location of multiplex-PCR primers

We used Minion, Oxford Nanopore Technology (ONT), to sequence the PCR product of Cemani with heterozygous allele due to poor sequence results from Sanger sequencing. We followed the protocol for the ligation sequencing kit (LSK 109) with the native barcoding kit (NBD104) with some modification. The raw fast5 data sequence was a base called and then demultiplexed using guppy (Guppy v 5.0.17) to separate samples based on their index. Fastq data was assembled using minimap 2.17 (Li, 2018) and then called the consensus sequence in Geneious Prime® 2021.1.1 (www.geneious.com). All sequence data generated by Sanger and Nanopore, ONT were then compared and analysed the polymorphisms in cutting position to identify the sequence pattern of each genotype. The diagram workflow of methods developed in this study is shown in Figure 3.

**Figure 3.**
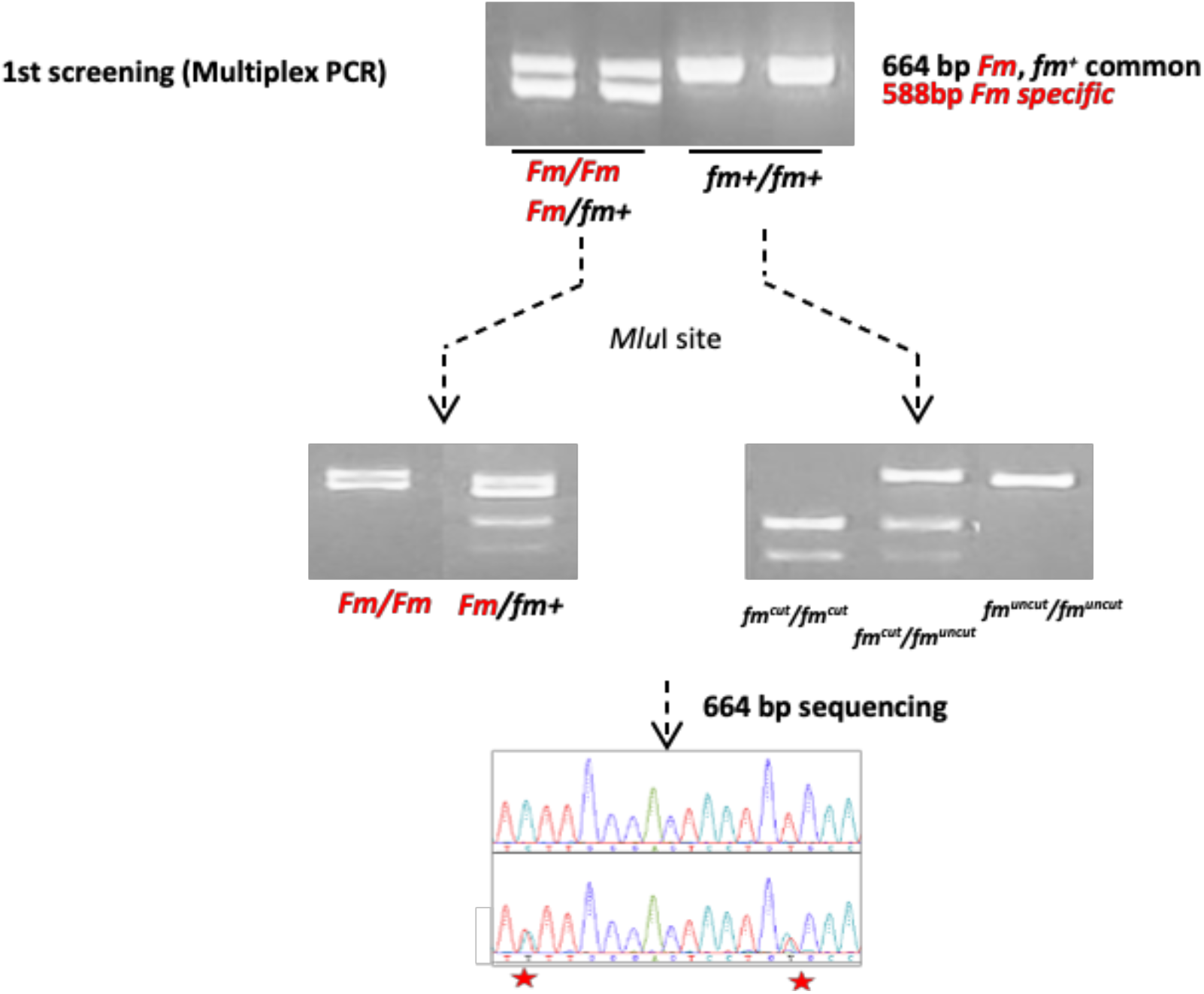
Workflow of PCR amplification, *Mlu*I restriction, and sequencing to characterize the *Fm* trait in Cemani **and non-*Fm*** chicken.

## 3. Results

### 3.1. The phenotype variation of the *Fm* trait in the Ayam Cemani population

Phenotypic information on Cemani chickens was obtained from the livestock research centre (Balitnak), Bogor and Sumber Unggas Indonesia (SUI). We examined the degree of blackness in the comb, skin colour (under wing) and cloaca of Cemani chicken lines in the two breeding centres. Two types of deep black and slightly light black in the melanin pigmentation in the comb, shank, skin underwing, and cloaca are shown in Figure 4. Then, the relationship between the individual differences in melanin pigmentation in the Ayam Cemani population and the Fm trait genotype was analysed by combining PCR and RFLP techniques, as described in Figure 3.

**Figure 4.**
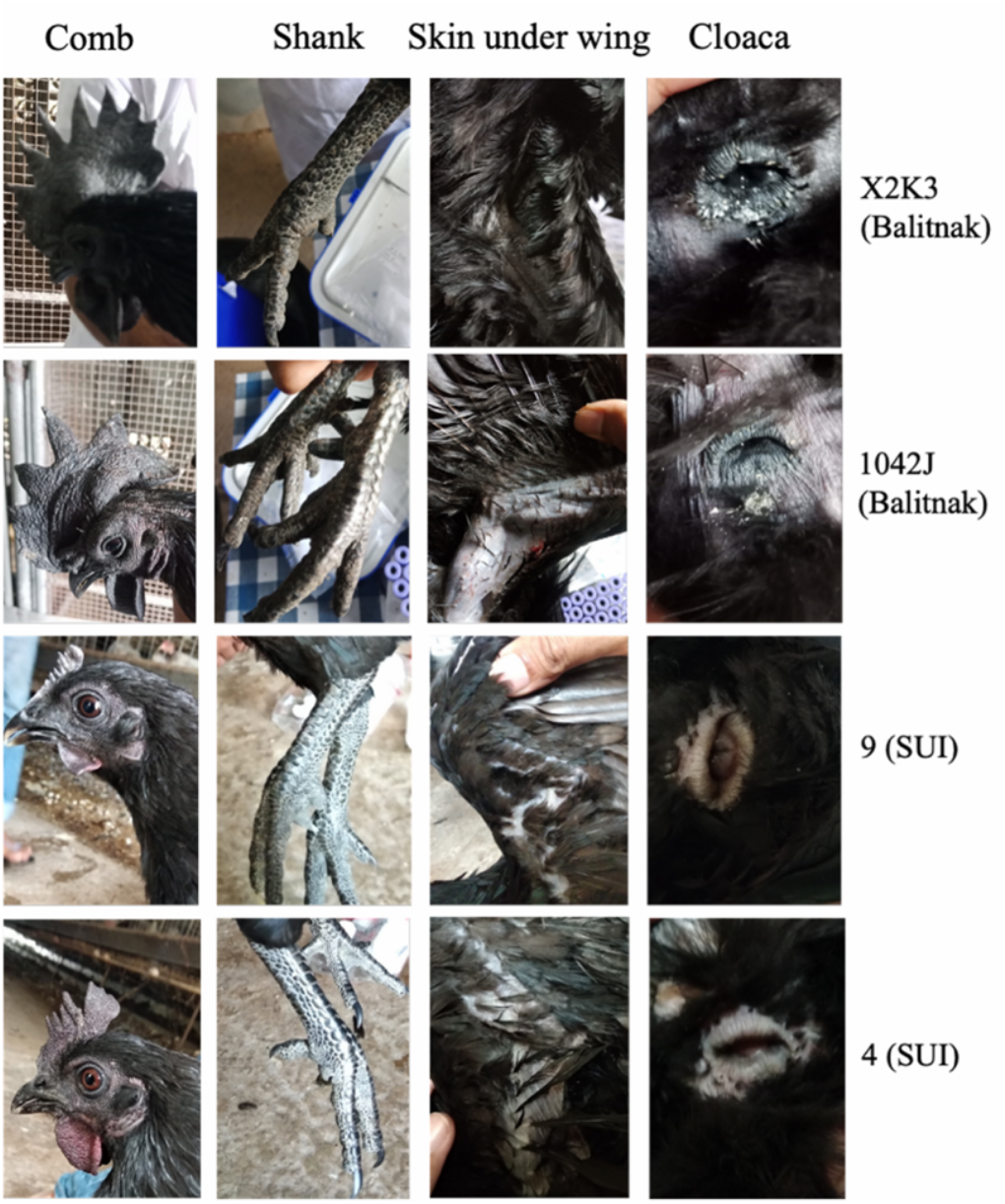
Color variation in the comb, shank, skin under wings, and cloaca of Cemani chicken.

### 3.2. Screening of *Fm* allele using PCR amplification and *Mlu*I RFLP in Cemani and non-*Fm* chicken population

For examining the variation in the duplication boundary of *Fm* in the Cemani chicken population, we used Cemani with the *Fm* mutation as a positive control (Cemani40), and we used IPB-D1, Kedu Putih (KDP), and broiler chicken (BROSS) as wild-type chicken without the *Fm* mutation. For *Fm* homozygous individuals, two copies of the 664 bp band common to the wild-type (WT, *fm*^+^) allele and two copies of the *Fm* allele-specific 588 bp band are amplified by PCR. However, for *fm*^+^ homozygotes (wild-type), only the 664 bp band is amplified. For heterozygous individuals, one copy of the 588 bp band (from the *Fm* allele) and two copies of the 664 bp band (from each of the *fm*^+^ and *Fm* alleles) are amplified. For differentiating *Fm* and *fm*^*+*^ homozygous and heterozygous genotypes, the presence or absence of the *Mlu*I restriction site within the 664 bp fragment is determined by PCR-RFLP or by sequence comparison of SNPs within the fragment.

*Fm*/*fm*^+^ heterozygotes as well as the *fm*^+^ alleles that have the *Mlu*I restriction site (*fm*^cut^) will produce three bands of 588, 400 and 264 bp length following *Mlu*I digestion. In contrast, *fm*^+^ alleles without *Mlu*I restriction site (*fm*^uncut^) will produce the undigested 664 bp fragment after RFLP. In case of *Fm*/*fm*^+^ heterozygotes with and without cutting position (*fm*^cut^/ *fm*^uncut^) will produce four bands of 664, 588, 400 and 264 bp length following *Mlu*I digestion. Therefore, wild-type chicken display variation in restriction patterns depending on their *fm*^+^ genotype. Wild-type *fm*^+^ chicken producing two bands (400 and 264 bp) after *Mlu*I digestion are *fm*^cut^/*fm*^cut^ homozygous, those producing three bands (664, 400, and 264 bp) after digestion are *fm*^cut^/*fm*^uncut^ heterozygous, and those producing an undigested 664 bp band following digestion are *fm*^uncut^/*fm*^uncut^ homozygous. The results of multiplex PCR amplification combined with RFLP using *Mlu*I restriction digestion are shown in Figure 5.

**Figure 5.**
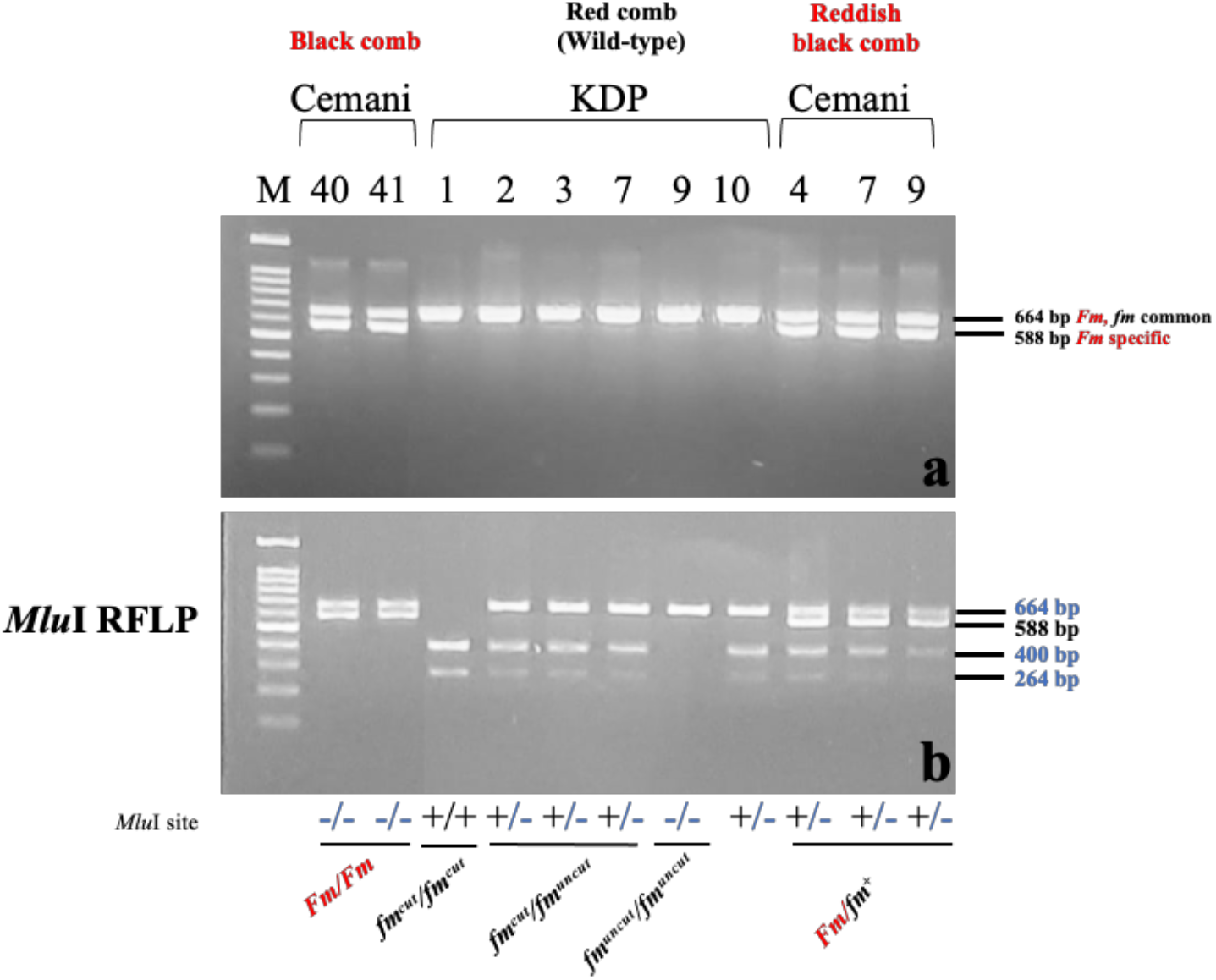
Gel electrophoresis of multiplex-PCR and RFLP products. Upper panel shows PCR products amplified by multiplex-PCR primers (a). Lower panel shows *Mlu*I-digested pattern of multiplex-PCR primer-amplified products (b)

In the first step of screening by PCR amplification using *Fm*-allele specific primers, the 588 bp fragment was amplified in 39 samples from Balitnak and 26 samples from SUI. For one Balitnak sample (355B) with the *Fm* mutation, the specific 588 bp fragment for the *Fm* allele was not amplified in the multiplex PCR, and the 664 bp fragment was not *Mlu*I-digested. This individual had the same phenotypic traits as other *Fm* chicken, and thus may have the *Fm*/*Fm* genotype, but some variation within the primer region may have resulted in failure to amplify the 588 bp fragment. From the total of 39 Cemani chickens confirmed to be *Fm*/*Fm* homozygous, 35 showed completely black phenotype in their combs, whereas four showed reddish black phenotypes in their combs and wattles (Table 2). Representative photos of two phenotypes among Cemani chickens with the homozygous *Fm*/*Fm* genotype (Cemani with either reddish black combs/wattles or black combs/wattles) are shown in Table 3. Unlike the RFLP results of the Cemani population in Balitnak, out of the 37 Cemani samples from SUI, 26 individuals were homozygous (*Fm*/*Fm*) and 11 were heterozygous (*Fm*/*fm*^+^) with four bands, whereas Cemani chicken have *fm*^+^ allele with and without cutting position (*fm*^cut^/*fm*^uncut^). (Figure 5: Cemani 4, 7, 9; Table 2). The previous study using Silkie chicken (*unreported study*) producing three bands (588, 400 and 264 bp) which mean Silkie chicken have *fm*^+^ allele with cutting position (*fm*^cut^/*fm*^cut^). Representative figures of Cemani with homozygous and heterozygous genotypes are shown in Table 3. Moreover, DNA samples from Cemani chickens collected in Kedu village revealed that all samples were *Fm*/*Fm* homozygous (Table 2). According to Sulandari (2007), these chickens showed black color in the skin, shank, tongue, and comb.

**Table 2.** Genotyping result of Cemani chicken in several sampling sites

| No | Cemani in Balitnak (n=40) |  |  |  |  | Cemani in SUI (n=37) |  |  |  |  | Cemani in Kedu village (n=29) |  |  |  |  |
| --- | --- | --- | --- | --- | --- | --- | --- | --- | --- | --- | --- | --- | --- | --- | --- |
|  | Sample ID | Fm phenotype | Sex | 664 sequence | Fm genotype | Sample ID | Fm phenotype | Sex | 664 sequence | Fm genotype | Sample ID | Sex | Fm phenotype | 664 sequence | Fm genotype |
| 1 | 84J | Reddish black | M |  | *Fm/Fm* | 1 | No picture |  |  | *Fm/Fm* | Cemani40 | F | Black | ✓ | *Fm/Fm* |
| 2 | 85J | Black | M |  | *Fm/Fm* | 2 | Black | F |  | *Fm/Fm* | Cemani41 | M | Black | ✓ | *Fm/Fm* |
| 3 | 355B | Black | F | ✓ | *Undetected* | 3 | Black | M |  | *Fm/Fm* | Cemani42 | M | Black |  | *Fm/Fm* |
| 4 | 389B | Black | F |  | *Fm/Fm* | 4 | Reddish black | F | ✓ | *Fm/fm<sup>cut</sup>* | CM2 |  | No picture |  | *Fm/Fm* |
| 5 | 1008B | Black | F |  | *Fm/Fm* | 5 | Black | F |  | *Fm/Fm* | CM4 |  | No picture |  | *Fm/Fm* |
| 6 | 1011J | Reddish black | M |  | *Fm/Fm* | 6 | Black | M |  | *Fm/Fm* | CM5 | M | Black |  | *Fm/Fm* |
| 7 | 1012B | Black | F |  | *Fm/Fm* | 7 | Reddish black | M | ✓ | *Fm/fm<sup>cut</sup>* | CM8 | M | Black |  | *Fm/Fm* |
| 8 | 1013J | Black | M |  | *Fm/Fm* | 8 | Black | M |  | *Fm/Fm* | CM9 | M | Black |  | *Fm/Fm* |
| 9 | 1014B | Black | F |  | *Fm/Fm* | 9 | Reddish black | M |  | *Fm/fm<sup>cut</sup>* | CM11 | M | Black |  | *Fm/Fm* |
| 10 | 1016B | Black | F |  | *Fm/Fm* | 10 | Black | M |  | *Fm/Fm* | CM12 | F | Black |  | *Fm/Fm* |
| 11 | 1017J | Reddish black | M | ✓ | *Fm/Fm* | 11 | Black | F |  | *Fm/Fm* | CM13 | F | Black |  | *Fm/Fm* |
| 12 | 1018J | Black | M |  | *Fm/Fm* | 12 | Black | F |  | *Fm/Fm* | CM14 | F | Black |  | *Fm/Fm* |
| 13 | 1019J | Black | M |  | *Fm/Fm* | 13 | Black | F | ✓ | *Fm/fm<sup>cut</sup>* | CM15 | M | Black |  | *Fm/Fm* |
| 14 | 1020J | Black | M |  | *Fm/Fm* | 14 | Reddish black | M | ✓ | *Fm/fm<sup>cut</sup>* | CM16 | F | Black |  | *Fm/Fm* |
| 15 | 1037J | Black | M |  | *Fm/Fm* | 15 | Black | F |  | *Fm/Fm* | CM17 | F | Black |  | *Fm/Fm* |
| 16 | 1038B | Black | F |  | *Fm/Fm* | 16 | Black | F |  | *Fm/Fm* | CM19 | F | Black |  | *Fm/Fm* |
| 17 | 1039B | Black | F |  | *Fm/Fm* | 17 | Black | F |  | *Fm/Fm* | CM20 | F | Black |  | *Fm/Fm* |
| 18 | 1041B | Black | F |  | *Fm/Fm* | 18 | Black | F |  | *Fm/fm<sup>cut</sup>* | CM21 | F | Black |  | *Fm/Fm* |
| 19 | 1042J | Black | M |  | *Fm/Fm* | 20 | Black | F |  | *Fm/Fm* | CM24 | F | Black |  | *Fm/Fm* |
| 20 | 1043J | Black | M |  | *Fm/Fm* | 21 | Black | F |  | *Fm/Fm* | CM25 | F | Black |  | *Fm/Fm* |
| 21 | 1044B | Black | F |  | *Fm/Fm* | 22 | Black | F |  | *Fm/fm<sup>cut</sup>* | CM26 | M | Black |  | *Fm/Fm* |
| 22 | 1045J | Black | M |  | *Fm/Fm* | 23 | Black | F |  | *Fm/Fm* | CM28 | M | Black |  | *Fm/Fm* |
| 23 | 1046B | Black | F |  | *Fm/Fm* | 24 | Black | M |  | *Fm/Fm* | CM29 | M | Black |  | *Fm/Fm* |
| 24 | 1047B | Black | F |  | *Fm/Fm* | 25 | Black | M |  | *Fm/Fm* | CM30 | M | Black |  | *Fm/Fm* |
| 25 | 1048B | Black | F |  | *Fm/Fm* | 26 | Black | M |  | *Fm/Fm* | CM31 | M | Black |  | *Fm/Fm* |
| 26 | 1049B | Black | F |  | *Fm/Fm* | 27 | Black | M |  | *Fm/fm<sup>cut</sup>* | CM33 | M | Black |  | *Fm/Fm* |
| 27 | 1050B | Black | F |  | *Fm/Fm* | 28 | Black | M |  | *Fm/Fm* |  |  |  |  |  |
| 28 | 1061B | Black | F |  | *Fm/Fm* | 29 | Black | M |  | *Fm/Fm* |  |  |  |  |  |
| 29 | 1063J | Black | M |  | *Fm/Fm* | 30 | Reddish black | M |  | *Fm/fm<sup>cut</sup>* |  |  |  |  |  |
| 30 | 1064J | Black | M |  | *Fm/Fm* | 31 | Reddish black | M |  | *Fm/fm<sup>cut</sup>* |  |  |  |  |  |
| 31 | 1065J | Black | M |  | *Fm/Fm* | 32 | Black | M |  | *Fm/Fm* |  |  |  |  |  |
| 32 | 1066J | Black | M |  | *Fm/Fm* | 33 | Black | M |  | *Fm/Fm* |  |  |  |  |  |
| 33 | 1068J | Black | M |  | *Fm/Fm* | 34 | Reddish black | M |  | *Fm/fm<sup>cut</sup>* |  |  |  |  |  |
| 34 | 1069J | Black | M |  | *Fm/Fm* | 35 | Black | M |  | *Fm/Fm* |  |  |  |  |  |
| 35 | 1070J | Black | M |  | *Fm/Fm* | 36 | Black | M |  | *Fm/Fm* |  |  |  |  |  |
| 36 | XK3B | Black | F |  | *Fm/Fm* | 37 | Black | M |  | *Fm/Fm* |  |  |  |  |  |
| 37 | X2K3J | Black | M |  | *Fm/Fm* | 38 | Black | M |  | *Fm/Fm* |  |  |  |  |  |
| 38 | X3K3B | Black | F |  | *Fm/Fm* |  |  |  |  |  |  |  |  |  |  |
| 39 | X4K3B | Black | F |  | *Fm/Fm* |  |  |  |  |  |  |  |  |  |  |
| 40 | X5K10J | Black | M |  | *Fm/Fm* |  |  |  |  |  |  |  |  |  |  |

Note : SUI : Sumber Unggas Indonesia (Breeding center), Balitnak: Livestock research center, Bogor.

**Table 3.**
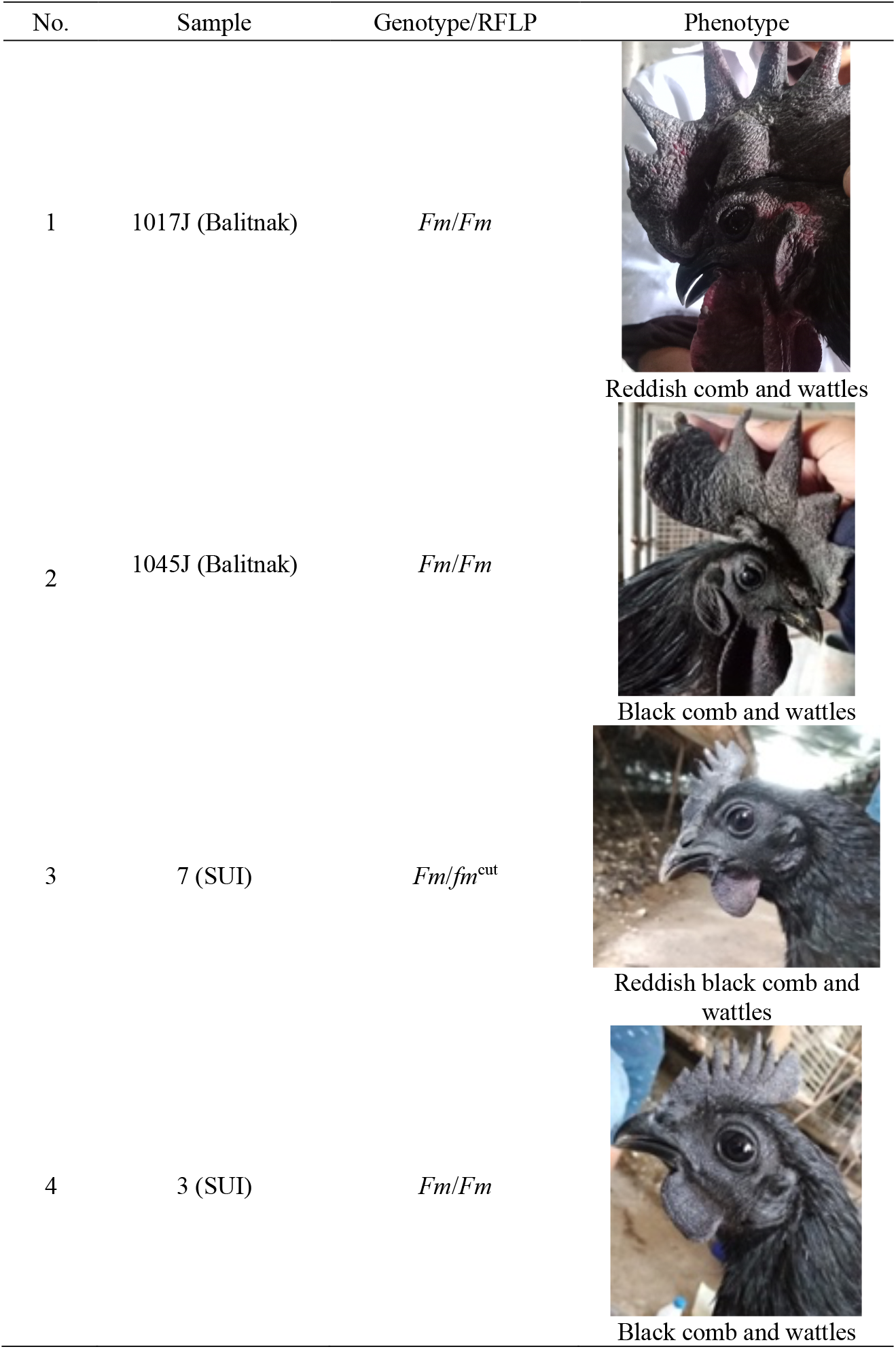
Representative of Cemani sample with phenotype and genotype No. Sample Genotype/RFLP Phenotype

| No. | Sample | Genotype/RFLP | Phenotype |
| --- | --- | --- | --- |
| 1 | 1017J (Balitnak) | *Fm/Fm* | Reddish comb and wattles |
| 2 | 1045J (Balitnak) | *Fm/Fm* | Black comb and wattles |
| 3 | 7 (SUI) | *Fm/fm<sup>cut</sup>* | Reddish black comb and wattles |
| 4 | 3 (SUI) | *Fm/Fm* | Black comb and wattles |

We also conducted MluI PCR-RFLP genotyping in 22 wild-type chickens with non-Fm phenotype such as Kedu Putih (KDP), Broiler, and IPB-D1 chickens. Multiplex PCR results showed that only the 664 bp fragment was amplified in all of these chickens. The 664 bp amplicons of some samples of KDP are shown in Figure 5a. After digestion of the 664 bp fragment with *Mlu*I, all Broiler chickens showed two bands, 400 and 264 bp in length. On the other hand, KDP chickens showed three genotypes (four individuals with *fm*^cut^/*fm*^cut^ [400 and 264 bp after digestion], three individuals with *fm*^cut^/*fm*^uncut^ [664, 400, and 264 bp after digestion], and one individual with *fm*^uncut^/*fm*^uncut^ [uncut 664 bp after digestion]) (Figure 5b, Table 4). In IPB-D1 chickens, two genotypes were detected (three individuals with *fm*^cut^/*fm*^cut^ and one individual with *fm*^cut^/*fm*^uncut^).

**Table 4.** RFLP result of Broiler, Kedu Putih, and IPB-D1 chicken using *Mlu*I

| No. | Broiler (n=9) |  | Kedu Putih (n=9) |  | IPB-D1 (n=4) |  |
| --- | --- | --- | --- | --- | --- | --- |
|  | Sample ID | Genotype/*MluI* site | Sample ID | Genotype/*MluI* site | Sample ID | Genotype/*MluI* site |
| 1 | BROSS1 | $fm^{cut}/fm^{cut}$ | KDP1 | $fm^{cut}/fm^{cut}$ | 9101 | $fm^{cut}/fm^{cut}$ |
| 2 | BROSS2 | $fm^{cut}/fm^{cut}$ | KDP2 | $fm^{cut}/fm^{uncut}$ | 294 | $fm^{cut}/fm^{cut}$ |
| 3 | BROSS3 | $fm^{cut}/fm^{cut}$ | KDP3 | $fm^{cut}/fm^{uncut}$ | 320 | $fm^{cut}/fm^{uncut}$ |
| 4 | BROSS4 | $fm^{cut}/fm^{cut}$ | KDP4 | $fm^{cut}/fm^{cut}$ | 381 | $fm^{cut}/fm^{cut}$ |
| 5 | BROSS5 | $fm^{cut}/fm^{cut}$ | KDP5 | $fm^{cut}/fm^{cut}$ | | |
| 6 | BROSS6 | $fm^{cut}/fm^{cut}$ | KDP7 | $fm^{cut}/fm^{uncut}$ | | |
| 7 | BROSS7 | $fm^{cut}/fm^{cut}$ | KDP8 | $fm^{cut}/fm^{cut}$ | | |
| 8 | BROSS8 | $fm^{cut}/fm^{cut}$ | KDP9 | $fm^{uncut}/fm^{uncut}$ | | |
| 9 | BROSS9 | $fm^{cut}/fm^{cut}$ | KDP10 | $fm^{cut}/fm^{uncut}$ | | |

### 3.3. Sequence data

To confirm the sequence polymorphisms within the 664 bp fragment including the *Mlu*I restriction site, each genotype was sequenced. The results revealed 18 SNPs between *Fm* and WT *fm*^*+*^ alleles. The *Mlu*I restriction enzyme recognizes the A^CGCGGT sequence, which is located at positions 401-406 of the 664 bp fragment and corresponds to positions 1239019-1239023 in the chicken genome assembly GRCg6a/galGal6 (Figure 6). The *Fm* homozygous birds (*Fm*/*Fm*) had T/T genotype at position 403 of 664 (galGal6, position 1239021) and produced an undigested 664 bp band following restriction analysis, whereas all wild-type *fm*^cut^/*fm*^cut^ birds had C/C genotype at position 403, and two bands of 400 and 264 bp were produced after digestion (Figure 6). Heterozygous Cemani posses cutting and uncutting position with T/C genotype produced four bands of 664, 588, 400, and 264 bp. However, in some wild-type birds without *Fm* mutation, substitutions from G to A were found at position 402 (galGal6, position 1239021) located within the *Mlu*I restriction site. The WT *fm*^uncut^ allele with A at position 402 was not digested by *Mlu*I, like for the *Fm* allele (Figure 6).

**Figure 6.**
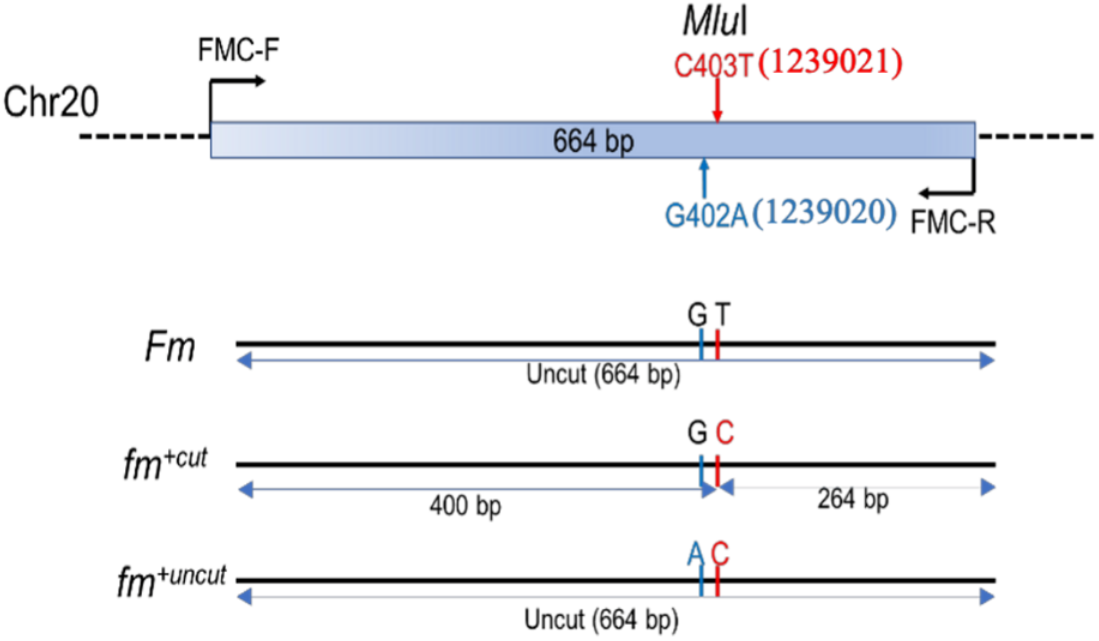
SNPs in *Mlu*I site within 664bp fragment in *Fm* allele and wild-type allele.

Each nucleotide was sequenced to confirm the sequence polymorphisms within 664 bp, including the MluI restriction site. The results revealed 18 SNPs within 664 bp between *Fm* and wild-type alleles, as predicted by *Mlu*I digestion of these two alleles. *Mlu*I restriction enzyme recognises A^CGCGGT sequences, which is at position 401-406 in sequence order of 664 bp corresponding to position 1239019-1239023 (chicken genome assembly GRCg6a/galGal6 (Figure 6). The *Fm* homozygous birds (*Fm*/*Fm*) had T/T genotype at position 403 (galGal6, position 1239021) of 664bp and produced an uncut band with 664bp, whereas the almost wild-type allele (*fm*^cut^/*fm*^cut^) had C/C genotype at position 403 and was digested into two bands of 400bp and 264bp (Figure 6). Thus, heterozygous Cemani (*Fm*/(*fm*^cut^/*fm*^uncut^)) with T/C genotype had four bands of 664, 588bp, 400bp, and 264bp from both alleles. However, in some wild-type birds without *Fm* mutation, substitutions from G to A were found at position 402 (galGal6, position 1239021) on the *Mlu*I recognition site. The wild-type allele (*fm*^uncut^) with A at position 402 was not cut by *Mlu*I and *the Fm* allele (Figure 6).

We found one SNP (C>T) within the *Mlu*I restriction site at position 403 and two insertions in the 664 bp fragment corresponding to position 1239021, 11239047 and 11239048 in chr 20; this SNP can distinguish three populations: *Fm* homozygous, *Fm* heterozygous, and wild-type chicken. There are a total of eight segregation sites between homozygous *Fm/Fm* and heterozygous *Fm/fm*^*+*^ chickens. There are no polymorphic sites within homozygous *Fm/Fm* chickens, whereas in the *Fm/fm*^*+*^ chicken population, most SNPs within the 664 bp fragment are heterozygous. One Cemani individual (355B) shared similar 664 bp sequence with *Fm* chickens, supporting that this individual is *Fm/Fm* homozygous. In the population of wild-type chickens, we found 15 segregation sites of which one SNP at position 11239020, located within the *Mlu*I restriction site, could differentiate the three wild-type genotypes: *fm*^*cut*^*/ fm*^*cut*^, *fm*^*cut*^*/ fm*^*uncut*^, *fm*^*uncut*^*/ fm*^*uncut*^ (Figure 7).

**Figure 7.**
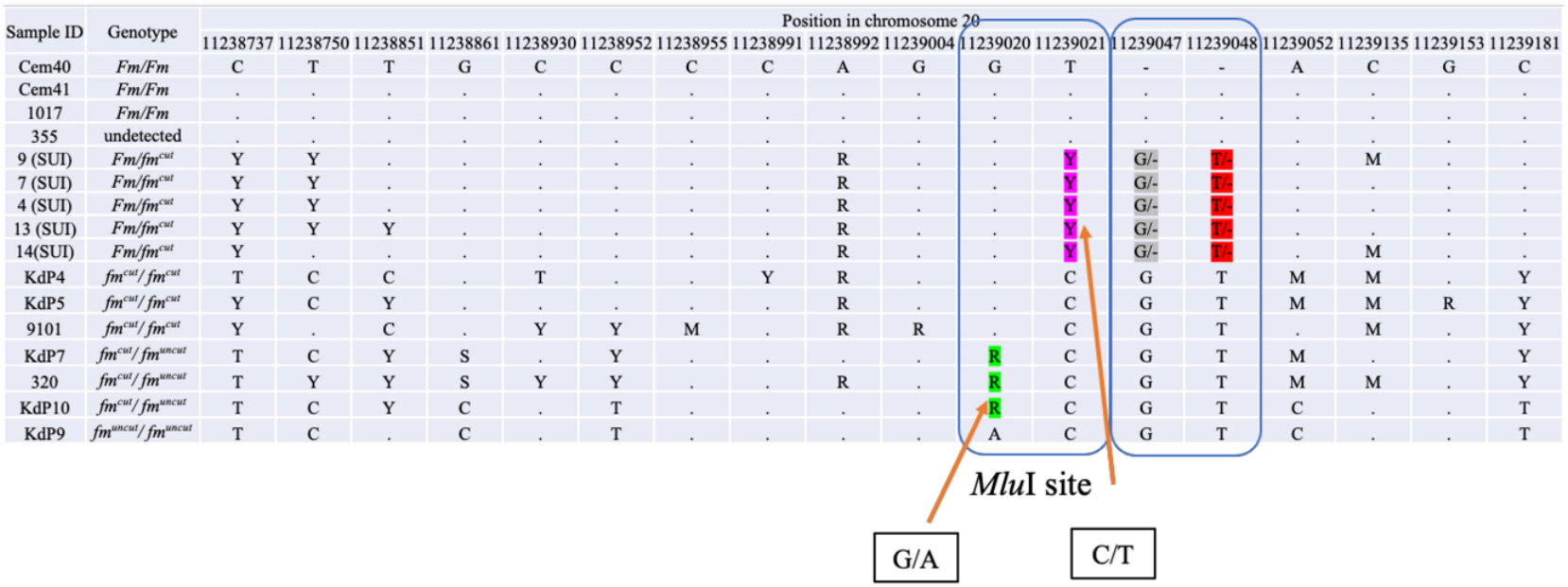
Single Nucleotide Polymorphism (SNP) in 664bp fragment in *Fm* and wild-type alleles.

## 4. Discussion

Previous molecular studies on skin pigmentation in domestic chicken include examining the causes of yellow skin color common in domestic chicken (Eriksson et al. 2008) and rare black colored skin caused by dermal melanocytes [i.e. Fibromelanosis (*Fm*)] (Dorshorst et al.2011, Shinomiya et al. 2011, Dharmayanthi et al. 2017). A recent genome-wide association study identified a genomic region on chromosome 20 including *EDN3* and *BMP7* that is associated with hyperpigmentation in chicken combs (Dong et al. 2019). In addition to the *Fm* locus, the dermal melanin inhibitor (*Id*) locus affects dermal hyperpigmentation (Dunn and Jull 1927). *Id* is important for controlling melanoblast migration routes, whereas *Fm* is responsible for melanoblast proliferation and maintenance, according to Dorshort et al. (2011).

In the present study, we applied PCR amplification followed by PCR-RFLP to reveal the different *Fm* genotypes in the Cemani population. PCR-RFLP is a simple technique that has been applied to species identification e.g. in bacterial (Vaneechoutte et al. 1996; Stubbs et al. 2000) or other contaminations in pork or chicken meat products (Erwanto et al. 2014; Djurkin et al. 2017). This method is also used for detecting variants related to traits such as growth and performance traits in chicken (Wang et al. 2017). The *Mlu*I PCR-RFLP method used in the current study was originally developed to determine homozygous or heterozygous birds of the *Fm* locus in Japanese Silkie (Ukkokei) with the *Fm* mutation, and using Black Minorca (non-*Fm*) chicken as a non-*Fm* reference breed. Applying this method in Cemani chicken was partly successful in determining the purity of Cemani chicken populations. The variation in blackness in the comb, shank, skin under wings, and cloaca of the present-day Cemani chickens in the breeding farm may be caused by crossbreeding between Cemani and other local chicken lines in the past. The *Fm* mutation is an autosomal dominant trait; thus, cross breeding with other chickens will produce progeny that have the *Fm*/*fm*^*+*^ genotype but are still hyperpigmented. Cemani with heterozygous traits may then become one of the founders in a breeding farm, resulting in the subsequent generation possessing either heterozygous or homozygous traits.

Most of the Cemani individuals with deep-black combs have the *Fm*/*Fm* homozygous genotype, whereas some individuals with reddish black combs possess the (*Fm*/(*fm*^cut^/*fm*^uncut^)) heterozygous genotype. Furthermore, some birds had brighter skin color and more reddish black combs. Based on RFLP results, we found that several of the Cemani chickens from Balitnak with reddish black comb were detected as *Fm* homozygous. However, some Cemani chicken with reddish black combs from SUI were *Fm* heterozygous. We propose one hypothesis that may explain this inconsistency. Since the individuals with reddish black combs have at least one copy of the *Fm* allele (confirmed by amplification of the 588 bp band), their genotype is either *Fm*/*Fm* or *Fm*/*fm*^+^. One limitation of our RFLP method is that *Fm*/*fm*^uncut^ chickens would be mistakenly identified as *Fm/Fm* chickens following *Mlu*I digestion of the 664 bp fragment, although *Fm*/*fm*^uncut^ chickens were not identified in our study. This limiation could be overcome by conducting both sequencing and RFLP analysis of the 664 bp fragment. Related to this, birds confirmed to be *Fm*/*Fm* genotype from sequence analysis but with wild-type phenotypes may be due to other factors. One possibility is that other pigmentation-related loci, such as the *Id* locus, suppress the expression of the *Fm* trait even though an individual may possess the *Fm* trait that is dominant over the wild type. The *Fm* trait appears as a dominant trait when mated with wild-type chickens, and generally, *Fm* homozygotes have a higher level of melanin deposition than *Fm/fm*^*+*^ heterozygotes. Therefore, the relationship between phenotype and genotype can be explained to some extent. In contrast, if there are other loci that suppress melanin deposition, this may decrease *Fm* expression, making it difficult to ascertain the genotype based on the phenotype. Therefore, genotyping at the molecular level is a suitable approach for solving such cases. Furthermore, in such cases, the internal tissues will most likely suppress pigmentation as well, making it difficult to distinguish based on the skin and comb. Conversely, if there are some factors that enhance the pigmentation, the situation may be even more complicated

Variation in the 664 bp fragment of the *fm*^*+*^ allele in WT chickens is also interesting, since these chickens have one of three genotypes: homozygous *fm*^cut^/*fm*^cut^, heterozygous *fm*^cut^/*fm*^uncut^, and homozygous *fm*^uncut^/*fm*^uncut^ genotypes. Heterozygous *fm*^cut^/*fm*^uncut^ indicates that one allele was not digested by *Mlu*I and homozygous *fm*^cut^/*fm*^uncut^ indicates both alleles were undigested. Variation in the *fm*^*+*^ allele may be due to polymorphisms at the *Mlu*I restriction site within the 664 bp fragment. Although there are 15 segregation sites within the 664 bp fragment, only one SNP distinguishes among the three genotypes in wildtype chicken. As mentioned above, if *Fm*/*fm*^uncut^ were to appear in the population, it would be difficult to distinguish from *Fm/Fm* using this method. To ensure the identification of pure *Fm* lines, there is an urgent need to develop RFLP based on other restriction sites within *fm*^+^.

Since we identified polymorphisms in the 664 bp sequence including two SNPs at the *Mlu*I site in *Fm* and WT chickens (Figure 7), *Mlu*I-RFLP and sequencing analysis are currently the best methods for conclusive confirmation of *Fm* genotypes. In addition, this method can clearly distinguish between homozygous *Fm*/*Fm* and heterozygous *Fm*/*fm*^*+*^ chickens (in Cemani the heterozygous written as *Fm*/(*fm*^cut^/*fm*^uncut^)). The PCR amplification, *Mlu*I-RFLP, and sequencing analysis of the causal region of the *Fm* mutation on chromosome 20 (developed in this study) were used to elucidate the state of fixation of the *Fm* trait in Cemani chicken. Furthermore, this combined molecular approach will be useful for determining the genotypes of the *Fm* mutation in Cemani chicken even in newly hatched chicks, for the purpose of selecting for pure Cemani-line with the *Fm*/*Fm* genotype. However, as the degree of blackness in outer tissues like comb or skin in Cemani chicken are probably affected by loci related to other melanin pigments as well as unidentified mutations, future determination of the *Fm-* related melanin inhibitors and/or activators in the comb and skin and their expression are necessary to fully elucidate the mechanism of hyperpigmentation in Cemani chicken.

## 5. Acknowledgements

Our thanks go to Shelvi, M. Fikri Al Habib (IPB University) and Rini Nuraeni (Research Centre for Biology (RCB)) for preparing the materials, and Dr. Quintin Lau for English editing and comments. This study supported by the *Skema Penelitian Dasar*, Ministry of Research and Technology/National Research and Innovation Agency, Indonesia, DIPA PN IPH-LIPI 2021, Avian Bioscience Research Center in Nagoya University, Japan, and the grant from SOKENDAI, Japan.

## 6. Author contribution

ABD design of the study and methodology carried out molecular laboratory work, contributed to data analysis and interpretation, drafted, reviewed and edited the manuscript. KK contributed to methodology and data interpretation and drafted, reviewed and edited the manuscript. IK carried out molecular laboratory work and helped draft the manuscript. RSM carried out molecular laboratory work and contributed to data analysis. SBI contributed to data analysis. Y and S carried out molecular laboratory work. ABLI and MSAZ contributed to the sample collection. YS and TA helped draft, review and edit the manuscript. CS designed the study and helped draft the manuscript. All authors gave their final approval for publication.

## Notes

### Competing Interest Statement

The authors have declared no competing interest.

